# Challenges in Using ctDNA to Achieve Early Detection of Cancer

**DOI:** 10.1101/237578

**Authors:** Imran S Haque, Olivier Elemento

## Abstract

Early detection of cancer is a significant unmet clinical need. Improved technical ability to detect circulating tumor-derived DNA (ctDNA) in the cell-free DNA (cfDNA) component of blood plasma via next-generation sequencing and established correlations between ctDNA load and tumor burden in cancer patients have spurred excitement about the possibilities of detecting cancer early by performing ctDNA mutation detection.

We reanalyze published data on the expected ctDNA allele fraction in early-stage cancer and the population statistics of cfDNA concentration to show that under conservative technical assumptions, high-sensitivity cancer detection by ctDNA mutation detection will require either more blood volume (150-300mL) than practical for a routine screen or variant filtering that may be impossible given our knowledge of cancer evolution, and will likely remain out of economic reach for routine population screening without multiple-order-of-magnitude decreases in sequencing cost. Instead, new approaches that integrate ctDNA mutations with multiple other blood-based analytes (such as exosomes, circulating tumor cells, ctDNA epigenetics, metabolites) as well as integration of these signals over time for each individual may be needed.

## Introduction

In the last decade, rapid technological development has dramatically improved our understanding of the underlying molecular basis of cancer [Vogelstein2013]. Such insights have contributed to the development of numerous molecularly targeted therapeutic agents and improved patient outcomes. Genomic tests are now used routinely to interrogate the genomes of patients with different diseases and identify targetable alterations. A frequent application of these tests is in cancer patients. Current estimates suggest that in the late-stage setting up to 50% of patients may receive genomic test results that alter their treatment [Blumenthal2016, LimaPereira2017]. However, the clinical utility of genomic tests remains controversial and is likely to evolve as new targets and their associated therapies emerge.

The finding that tumor-derived genomic alterations are detectable in “cell-free DNA” circulating in the plasma of patients with malignancy [Sorenson1994, Nawroz1996] has inspired the development of blood-based assays for tumor genomic profiling. In particular, sequencing of circulating cell-free tumor DNA (ctDNA) has been employed as an adjunct to DNA derived from tissue biopsies to inform treatment decision-making with the first such test (Guardant360, Guardant Health) launched in 2014 [Lanman2015]. ctDNA can be used to monitor minimal residual disease following treatment [GarciaMurillas2015, Chaudhuri2017], as well as the emergence of resistant clones prior to clinical resistance detectable via imaging [Misale2012, Diaz2012].

Not surprisingly, there is significant interest in the potential utility of ctDNA for the early non-invasive detection of cancer [Table 1], with over 1 billion USD invested in companies developing such technologies in 2017 alone [CNBC2017, GlobeNewsWire2017]. Indeed, ctDNA is detectable in some patients with early-stage cancers [Bettegowda2014, Phallen2017], but assay sensitivity has so far precluded their use for screening. The motivation for early detection of tumors is clear. It has long been appreciated that earlier detection of malignancy results in significant reduction in cancer-specific mortality [Etzioni2003, Cho2014]. This has led to screening guidelines for breast cancer via mammography and colorectal cancer via colonoscopy or stool-based assays, but such approaches are cancer-type specific. Serum protein biomarkers such as carcinoembryonic antigen (CEA) and cancer antigen 125 (CA-125) are often used to monitor disease progression; however, generalizable biomarkers that detect cancer early have remained elusive.

**Table 1:**
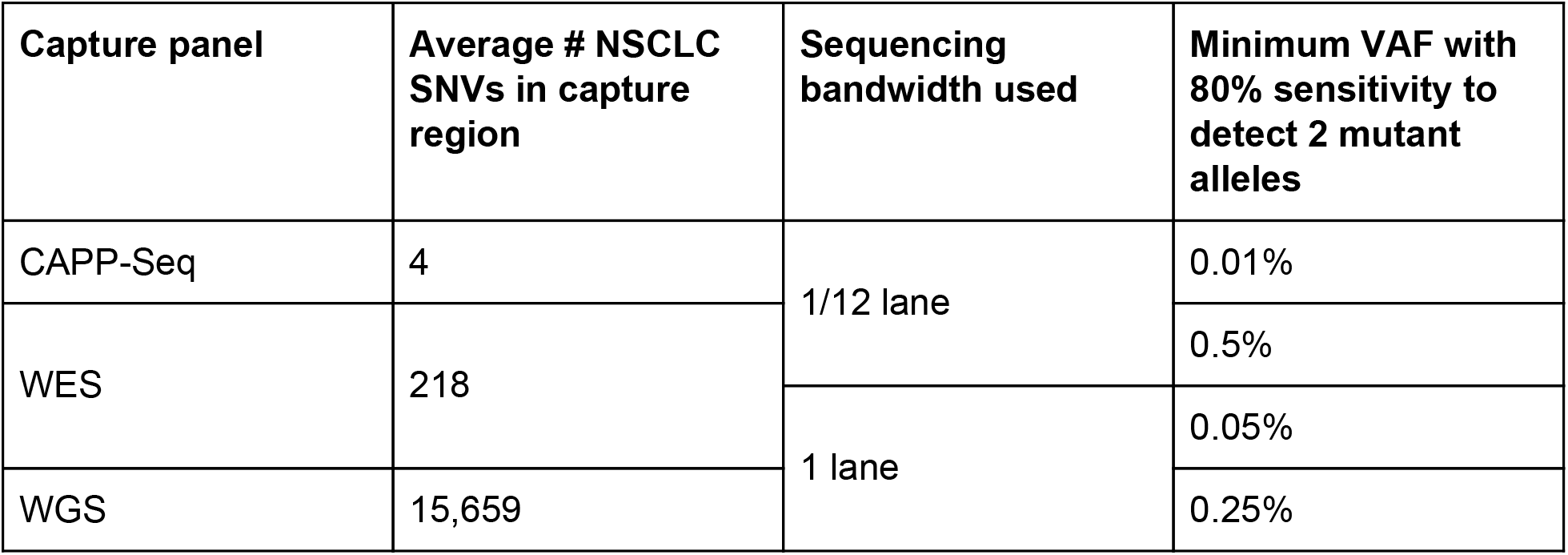
Effect of panel size and sequencing bandwidth on ability to detect somatically mutated alleles in plasma. Data from [Newman2014], figure 1d.

Aravanis and colleagues [Aravanis2017] recently proposed that the fundamental limitations for such a ctDNA-based early detection test, beyond the current state-of-the-art, are a requirement for around 100x more sequencing bandwidth and improved variant interpretation. Subsequently, in the most comprehensive study of early stage cancers to date, Phallen et al. [Phallen2017] reported the detection of somatic alterations in 50-75% of patients depending on histology. While this represents a significant advance over prior reports, achieved through both deeper panel sequencing and improved variant calling via error correction, the sensitivity remains well below that required for broad clinical implementation. Despite the obvious shortfall of this and prior approaches, there has been limited examination of the feasibility of improving sensitivity to the levels required to demonstrate clinical utility. Here, we reanalyze published ctDNA sequencing data from early-stage cancer patients and assert that statistical and physiological limitations suggest that a ctDNA-based mutational assay for early detection would be neither commercially nor biologically viable.

## Targeted ctDNA mutation-detection panels require infeasibly large input volumes for early detection

Detection of tumor-derived alleles in the blood can be modeled as a binomial process: we sequence a number of independent fragments of DNA, of which only a fraction (given by the variant allele fraction, or proportion of cfDNA at a given locus carrying a tumor-derived mutation) will be derived from the tumor. In order to detect cancer, the most sensitive test that could be built would be one that reported “positive” upon the detection of a single fragment carrying a cancer-derived allele. However, tumors exhibit a remarkable degree of mutational heterogeneity and the initiating lesions are largely unknown. This is further complicated by the presence of somatic alterations in normal tissue [Martincorena2015] and blood cells (so called clonal hematopoiesis of indeterminate potential, or CHIP) [Jaiswal2014, Genovese2014, Xie2014]. Although in practice such factors will contribute to an unacceptably high false-positive rate, this simple model sets a *lower* bound on the amount of sequencing required. If such a test were required to be 95% sensitive, this would be equivalent to requiring that in 95% of samples that carry a tumor, at least one fragment would be detected by the test. **Figure 1a** illustrates the sensitivity of a test modeled by this process, as a function of variant allele frequency (VAF) and (unique) sequencing depth.

**Figure 1:**
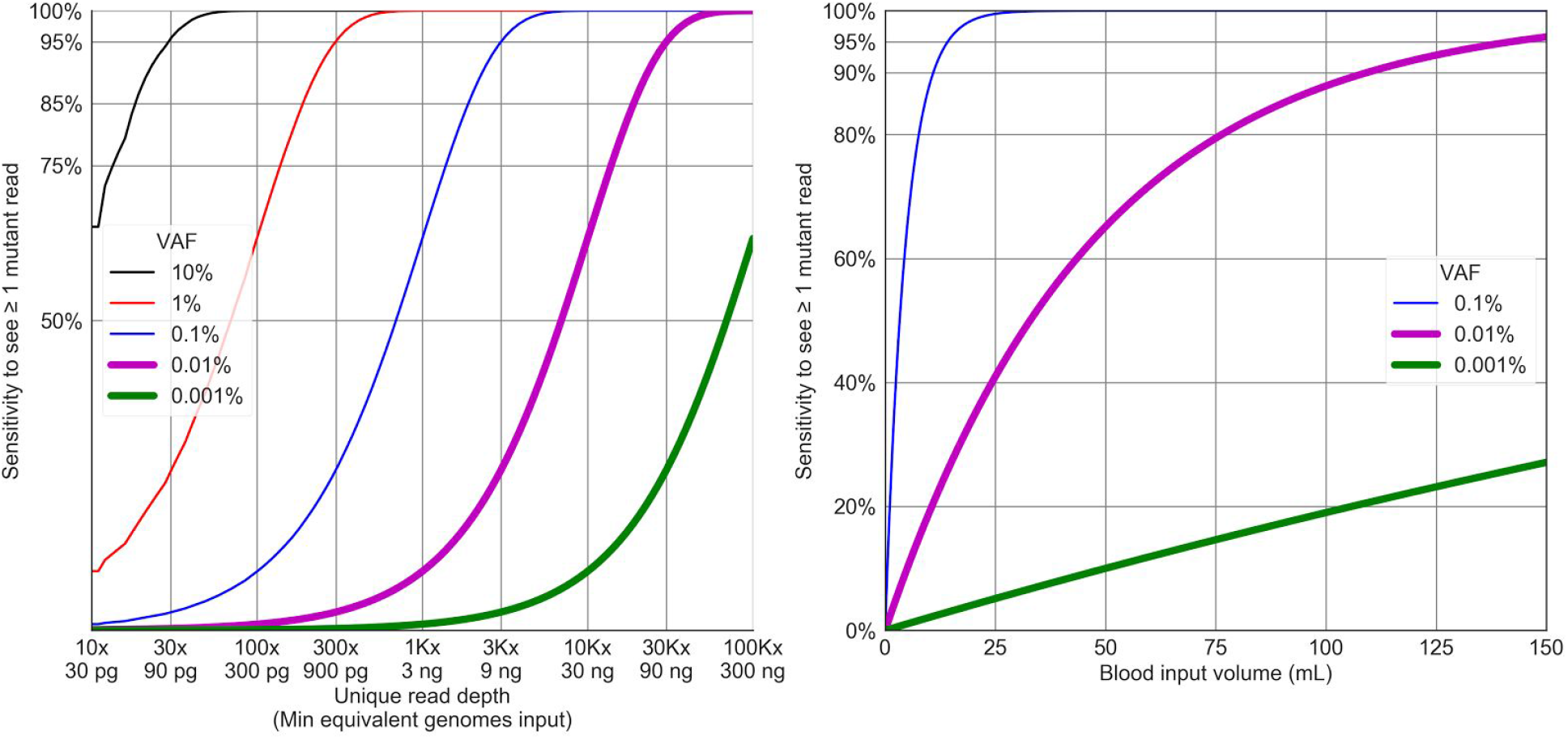
Binomial model for ctDNA sequencing: sensitivity achieved by a sequencing assay to detect 1 mutant allele at a given VAF. **a)** Upper bound on sensitivity as a function of sequencing depth; note logarithmic x-axis. Also shown is the minimum amount of unique DNA input required for sequencing, assuming 3pg haploid genome mass and 100% process efficiency. **b)** Sensitivity as a function of blood input volume, assuming 2.3ng cfDNA/mL plasma, plasma volume 55% of blood volume, and 50% process efficiency.

Aravanis and colleagues have suggested a VAF target for early detection of 0.01% [Aravanis2017] based on clinical data indicating that in early-stage cancer patients the fraction of cfDNA attributable to ctDNA (“tumor fraction” or “ctDNA fraction”) is on or below this order of magnitude. Bettegowda and colleagues were able to detect ctDNA in 47-55% of 182 stage I and II cancer cases [Bettegowda2014]. Newman et al. sequenced plasma from 13 patients with non-small-cell lung cancer (NSCLC) and found that 63% of 11 patients exhibited ctDNA fraction ≤0.5%, including all four patients with stage I NSCLC; more generally, the ctDNA fraction rose with tumor volume [Newman2014]. More recently, Phallen and colleagues sequenced an 81-kb region in cfDNA to an average unique coverage of 6,182X in 138 stage I/II solid tumor patients [Phallen2017]. **Figure 2a** illustrates the distribution of tumor VAF observed in their stage I/II patients: 50% of stage I and nearly 30% of stage II patients had no mutated ctDNA observed. In the remainder, most patients had maximal ctDNA VAFs between 0.1% and 1%; however, a steep cutoff is visible in the plot below 0.1%.

**Figure 2:**
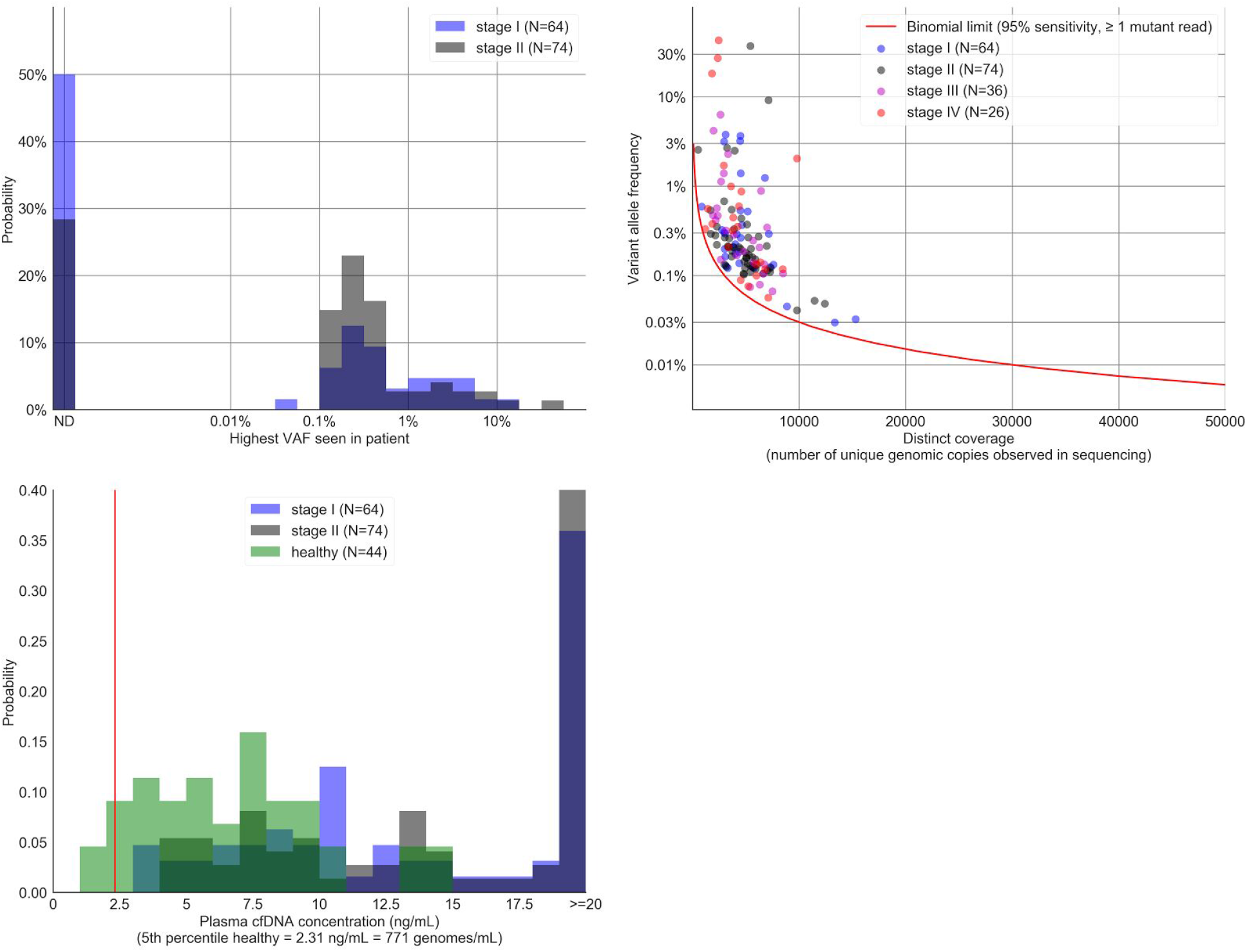
Analysis of ctDNA characteristics from early-stage cancer patients in Phallen et al. [Phallen2017]. Panels clockwise from top left. **a)** Highest per-patient VAF observed for any cancer-related variant in stage I/II colorectal, breast, ovarian, or lung cancer patients, as measured by the TEC-Seq protocol. Samples listed as “ND” had no cancer-derived alleles observed. In patients with multiple cancer alleles detected in plasma, the highest VAF is shown. **b)** Comparison of observed VAF to binomial model. Dots correspond to the VAF of the lowest-frequency cancer-derived variant detected and the unique depth of coverage for that patient. Red and blue curves respectively show the VAF expected to be detected with 95% confidence by the binomial model in Figure 1. **c)** Distribution of cfDNA concentrations observed in healthy individuals and stage I/II cancer patients. Red line is drawn at the 5th percentile of the distribution of healthy individual cfDNA concentration.

The binomial model for ctDNA detection suggests that the cutoff visible in **Figure 2a** may arise from the underlying assay having insufficient depth to recover rarer alleles. **Figure 2b** shows patients sequenced by Phallen et al. by their *lowest* VAF allele, as a function of the unique coverage achieved on each patient. Also illustrated is the 95% sensitivity bounds derived from the binomial model in Figure 1a; if the models held, we would expect that these lines would represent the lower bounds on the data. Indeed, this is observed: no patients appear below the 95% sensitivity limit, and the handful of patients with alleles detected at VAF below 0.05% also have higher sequencing depth, tracking with the model predictions. Thus, published data suggests that a substantial fraction of patients (perhaps 50% of stage I cancer patients) have tumor-derived alleles present below 0.1% that have not been sequenced deeply enough to detect. This model thus suggests a unique coverage of 30,000x as a target for an early detection test (yielding 95% sensitivity for one read of a single allele at 0.01%).

We have so far assumed that it is possible to arbitrarily increase sequencing depth as required to detect rare alleles. Also illustrated in **Figure 1a** is the genomic input required for the test to achieve this level of unique coverage (assuming 100% process efficiency from DNA extraction through sequencing): it is not possible to have 30,000 unique reads without 30,000 unique copies of the locus being sequenced. **Figure 2c** illustrates the amount of cfDNA that can be expected from healthy individuals and those with early stage cancer, drawing from data in [Phallen2017]. To interpret these data, it is critical to consider the context in which an early detection test would be run: in a mostly healthy population, as a test with an expectation of having a result delivered. Consequently, it is appropriate to consider summary statistics of the healthy distribution; since cancers are rare, most patients will be healthy. Furthermore, the mean and median are not the most useful measurements: if a test failure rate less than 5% (i.e., that 95% of prescribed tests successfully return a result) is required, this implies that the minimum amount of DNA acceptable for the test can be no higher than the 5th percentile of the population distribution. This is illustrated in **Figure 2c**: although the mean healthy individual has a plasma cfDNA concentration of 6.6 ng/mL, an individual at the 5th percentile has 2.3 ng/mL, and it is this concentration that determines the practical limitation for assay sensitivity.

Based on the minimum input established in **Figure 1a** by sampling considerations, and the population distribution from **Figure 2c**, we can now compute the total amount of plasma or blood input required for an early detection test using this approach: assuming 100% process efficiency (every molecule of cfDNA in the collection tube makes it to the sequencer), 90 ng of cfDNA are required for 95% sensitivity to detect 0.01% VAF alleles. We can expect at least 2.3 ng cfDNA/mL of plasma and therefore must collect at least 39.1 mL of plasma from each patient; since plasma constitutes ~55% of total blood volume, we must collect at least 71.1 mL of blood per patient. However, 100% process efficiency is unrealistic: sequencing involves a number of lossy steps (extraction, library preparation, target capture, sequencer loading, etc.). **Figure 1b** illustrates the constraint imposed by cfDNA concentration and process efficiency, modeling sensitivity at various VAF levels as a function of blood input volume and assuming 50% process efficiency. Even high total process efficiencies of 25-50% imply a total blood collection volume of 150-300 mL -- a substantial fraction of the approximately 5000 mL of total blood in an adult!

Thus, analysis of a binomial model that assumes constant VAF suggests that detecting alleles at 0.01% VAF with 95% confidence would require 150-300mL blood collection with 30,000x *unique* depth of sequencing coverage. Even if such amounts can in theory be collected over repeat blood draws, the logistics and limits of human physiology make the approach impractical for a population screening test. Alternative approaches that reduce the input requirements are therefore required.

## Broad mutation-detection panels require infeasibly high sequencing bandwidth and are limited by somatic heterogeneity

Data from The Cancer Genome Atlas (TCGA) and other large-scale cancer genomics projects have revealed that most tumors harbor hundreds to thousands of somatic variants [Kandoth2013], a subset of which are are highly recurrent across patients and tumor types [Ciriello2013]. This affords an opportunity to improve on the binomial sampling bounds derived in the previous section: If it is acceptable to detect *any* of a large number of tumor-derived mutations, instead of one or a few mutations, then for the purposes of the binomial model the effective VAF becomes less than or equal to the sum of the individual VAFs (with equality achieved if the presence of each variant is independent of the others). That is, to detect any of ten independent VAF=0.01% mutations has the same sampling difficulty as detecting a single 0.1% variant. Because this strategy is able to extract information from multiple loci, it gets more out of each genome, reducing input requirements as well as depth requirements.

In the best case, if all mutations had the same frequency and all were independent, a panel capturing N potential variant loci would have N times the power (and require 1/N the genomic input) of a single-site assay. However, real-world data from tumor sequencing suggests that this best-case scenario is not likely to be achieved in practice, because mutation frequency distributions are not uniform. This then requires additional sequencing bandwidth. Newman and colleagues developed a protocol (CAPP-Seq) to sequence recurrently mutated regions in NSCLC and modeled its analytical performance versus whole-exome and whole-genome sequencing. **Table 1** shows the results of their model: more extensive panels (e.g., whole-genome vs CAPP-seq) were able to capture more mutations, but this actually reduced sensitivity (even when given additional sequencing bandwidth). Thus, although panel expansion may offer a way to reduce input requirements the non-uniform distribution of mutation frequencies implies that dramatically more sequencing depth is required to recover comparable sensitivity to assays focused on high-frequency sites.

Furthermore, a well understood problem in next-generation sequencing (NGS) is that per-base error rates in NGS reads are substantially higher than 0.01%, often in the 0.1-0.5% range [Minoche2011]. A variety of techniques utilizing molecular barcoding and computational postprocessing have been developed to reduce this error rate [Schmitt2012, Newman2016, Phallen2017] but with the tradeoff of requiring higher read depth: individual molecules are sequenced more than once, with unique molecular identifiers used to group multiple reads of a single read and correct errors by consensus. This oversampling therefore inflates the total sequencing depth (as opposed to unique sequencing) required to perform a ctDNA assay: in TEC-Seq [Phallen2017], the reported interquartile range of oversampling was 4.77- to 9.38-fold, with a mean of 9-fold. Therefore, error correction may conservatively increase sequencing requirements fivefold.

An assay targeting an ideal (independent, uniform frequency) panel of mutations with cumulative VAF of 0.01% would require 30,000x unique coverage or, optimistically 150,000x raw depth with error correction. However, as established in the previous section, such a panel would require too much blood to be feasible. An ideal panel with 0.1% VAF might only require 3,000x unique coverage, and therefore 15-30mL blood collection; however, the CAPP-Seq experience suggests that in fact the total amount of sequencing required may be tenfold higher than that of the small panel -- equivalent to 1,500,000x coverage of the smaller panel! Thus, while panel expansion may constrain input requirements, it can only do so at an exorbitant sequencing cost.

Beyond mere technical difficulty, expanding the region of interest poses a fundamental biological challenge. Recent data exploring the mutational landscape in healthy individuals has revealed mutations in cancer driver genes at frequencies comparable to the VAF ranges being explored in early detection tests. For example, Martincorena and colleagues assayed healthy sun-exposed eyelid skin and found a dense landscape of low-level somatic variants in cancer genes such as *NOTCH1/2/3, TP53, FGFR3, FAT1,* and *RBM10* [Martincorena2015]. Clonal hematopoiesis associated with aging [Jaiswal2014, Genovese2014, Xie2014] also represents a major confounder of mutational heterogeneity in ctDNA. Results summarized in **Table 2** show that 10-40% of individuals carry low-level somatic mosaicism in cancer-associated genes. The presence of such variants in healthy individuals is thus a significant challenge for early cancer detection by mutation analysis because many such variants are expected to be present in the blood of older individuals (the intended population for cancer screening) and since hematopoietic stem cells dominate cfDNA signals [Snyder2016]. Moreover, the binomial limits derived above are highly optimistic - many of the detected alleles may need to be filtered out and more than one mutation will likely be required to mitigate false positives.

**Table 2:**
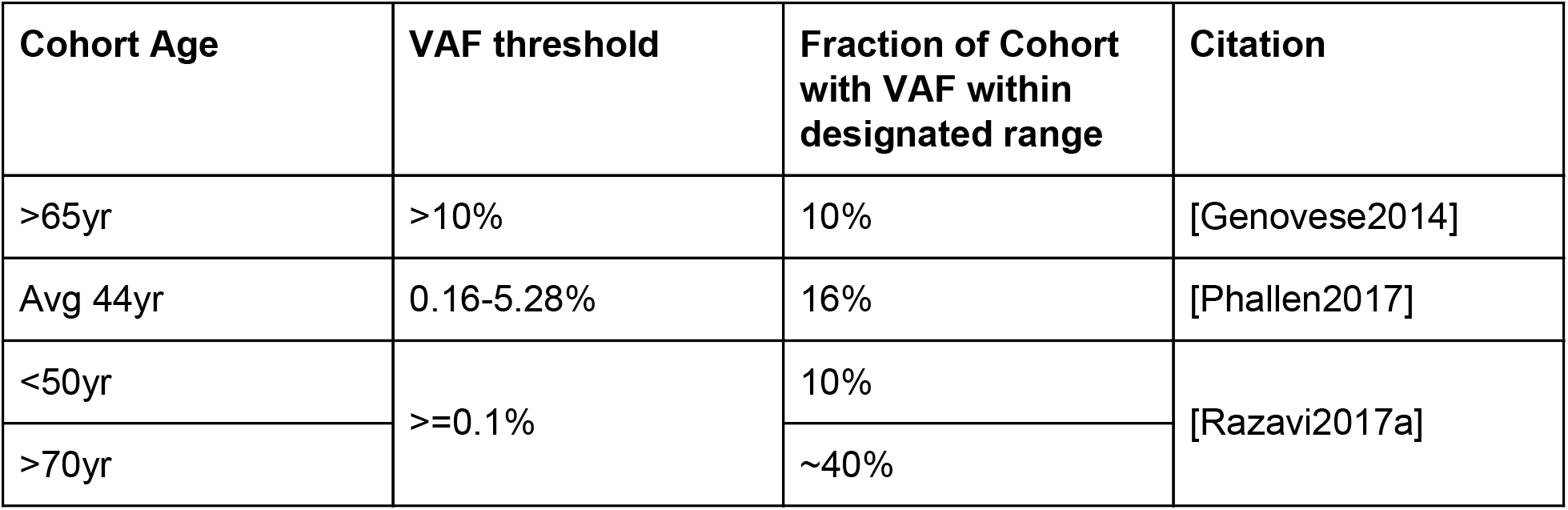
Prevalence of somatic variation observed in genes associated with clonal hematopoiesis.

It is increasingly appreciated that many somatic variants are present prior to transformation and malignant outgrowth, reflective of the relatively long time period leading up to cancer formation [Tomasetti2013, Sottoriva2015]. The presence of such variants is likely to complicate the task of identifying the presence of a tumor. As the spectrum of somatic variation in a given healthy organ tends to resemble the spectrum found in tumors originating in that tissue [Hoang2016], it may be difficult to distinguish which mutations are truly indicative of a tumor’s presence, and a difference in the rate at which tumor cells and the surrounding normal tissue shed cfDNA would be needed to extract signal [Hori2011].

Tumor initiating mutations will be present in all cells of the tumor (clonal), and therefore should be more readily detectable in ctDNA than subclonal mutations would be [Izumchenko2015], and may be more specific than passenger mutations that may occur in the peripheral normal. Indeed, established tumor initiating mutations and presumed cancer driver genes that are recurrently mutated across patient tumors have formed the basis for mutational-based ctDNA assays. However, even when such canonical driver mutations (*KRAS, PIK3CA, BRAF*) are present in the tumor tissue, they may not be detectable in corresponding ctDNA using the most sensitive techniques (ddPCR or TEC-seq) [Phallen2017], thus highlighting the challenge of mutation-only based strategies. While the landscape of somatic alterations in solid tumors have been catalogued, defining the *functional* drivers of individual cancers remains a challenge due to the high degree of mutational heterogeneity and the fact that drivers are both cell-of-origin and context dependent. Further, it is well established that multiple alterations are necessary for tumor development and that they need not be exclusively genomic [Vogelstein2013]. As such, biomarkers of malignant transformation in diverse histologies remain elusive and a better understanding of the earliest events of tumorigenesis could inform early detection efforts.

## Health Economics of Mutation Calling for Early Detection

Early screening tests for cancer are not novel to the medical system; existing tests provide a benchmark for the costs that payers are willing to tolerate for screening in the general population. As an example, in the USA, Medicare agreed to reimburse the Cologuard fecal screening test for colorectal cancer at $502 every three years [Pickhardt2016]. Assuming the physical limitation of sample volume and interpretive limitations of somatic heterogeneity could be successfully overcome, reimbursement may still pose a fundamental economic threat to the viability of mutation-based ctDNA assays for early detection. This is likely to be true even for a broad test intended to screen for most common solid cancers, for which the appropriate benchmark would be the *combined* costs from each individual traditional screen.

It is possible to estimate the costs of running a mutation detection assay using simple assumptions about test parameters. **Table 3** estimates the input volume required and sequencing cost of a mutation-calling ctDNA-based early detection assay under highly conservative assumptions. Under these assumptions, it is evident that reasonably sized panels for tumor liquid biopsy are commercially possible with present-day technology: 15 mL sample draw (compared to 20 mL reported in a real-world validation [Lanman2015]), with sequencing cost in the tens to hundreds of USD. However, early detection appears infeasible: while small panels (e.g., the 81-kb TEC-Seq panel) have achievable sequencing costs, their input volumes are likely prohibitive. In contrast, larger panels (e.g., the 2-Mb panel reported by Razavi and colleagues [Razavi2017b]) have sequencing costs alone that are nearly tenfold the total reimbursement for existing screening tests -- even without considering the likely increase in sequencing required to exploit lower-VAF mutations in the expanded panel or the need to perform repeat assays to improve assay sensitivity (or evaluate a change in ctDNA abundance). While sequencing costs fell dramatically in the early days, in recent years such cost reductions have slowed dramatically, and technological advances have focused on physical or synthetic long-read technologies that do not aid sequencing of short (~170 bp) cfDNA fragments. Consequently, it is not clear that the large cost gap between value-based reimbursement and sequencing cost for ultra-deep ctDNA mutation calling could be bridged in the near term.

**Table 3:**
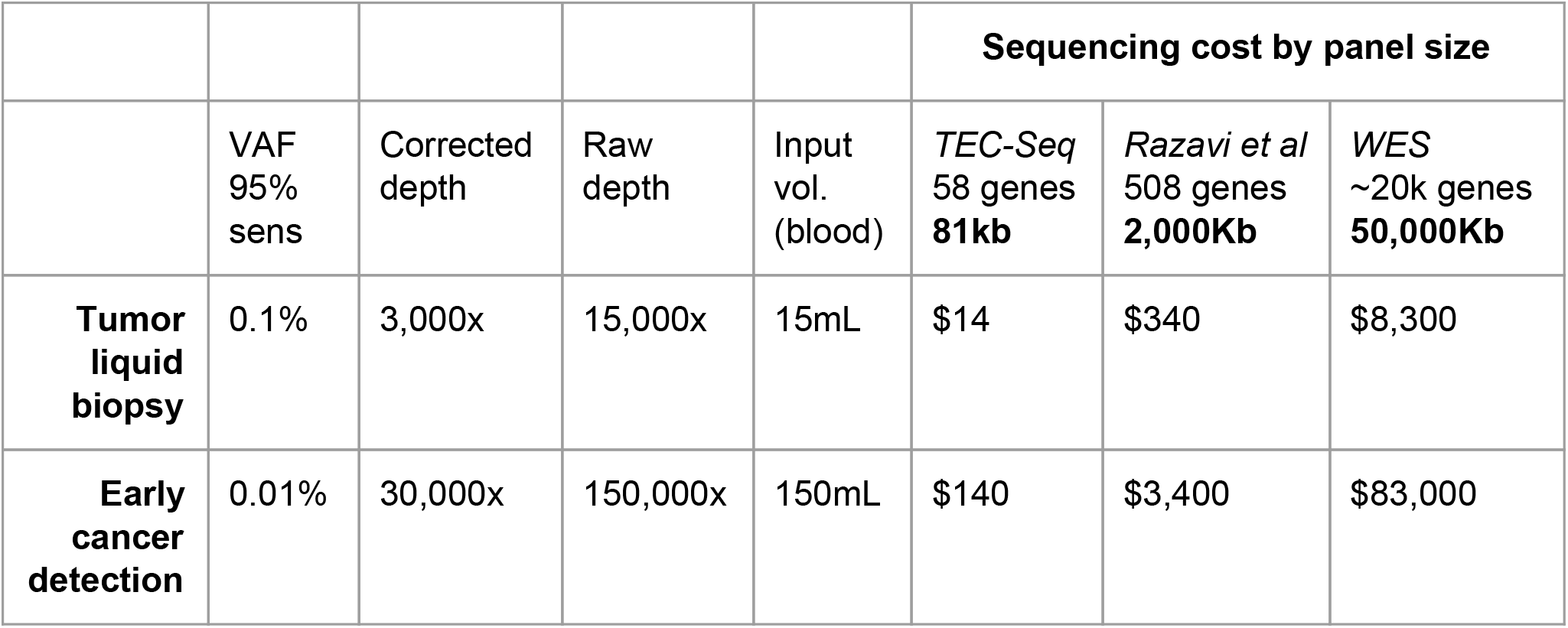
Assay requirements for tumor liquid biopsy and mutation-based early cancer detection. Assumptions as described in the main text:

- No more than 5% of samples may fail because of insufficient cfDNA quantity.
- We require 95% sensitivity to detect one read from any cancer-derived allele, assuming that one is present in the sample.
- 50% process efficiency: half of the cfDNA molecules in the input blood sample are represented in the sequencer output.
- 5x oversampling in sequencing for error correction.
- 100% on-target rate in target enrichment.
- “$1000 genome” sequencing costs: US$1000 / (30 × 3 Gbp) of sequencing bandwidth
- Only sequencing costs computed; all other costs (labor, equipment, facilities, depreciation, etc.) accounted at $0.
- Panel expansion neither reduces input requirements nor increases sequencing requirements.

## Summary: Clinical Performance of Early Detection by Mutation Calling

Although the challenges outlined in the previous sections are formidable, if it were possible to overcome them to significantly improve on current screening efforts, there may yet be value in pursuing this approach. However, clinical data (summarized in **Table 4**) casts doubt on this. Phallen and colleagues analyzed 138 individuals with stage I/II breast, colorectal, lung, or ovarian cancer using the TEC-Seq ctDNA 81-kb panel, at an average effective (unique, error-corrected) coverage of 6,182x [Phallen2017]. This method achieved a clinical sensitivity (fraction of clinically affected patients called positive by the assay) of 59-71% depending on disease stage, with tumor fraction and sequencing coverage the probable limiting factors.

**Table 4:** Comparison of ctDNA-based cancer detection assays versus conventional screening tests.

| ctDNA tests |  |  |  |  |  |  |
| --- | --- | --- | --- | --- | --- | --- |
| Cancer type | Stage | Total Samples | Panel size | Avg Raw Depth | Avg Eff Depth | Clin. Sens. |
| Breast, Lung, Colorectal, Ovarian [Phallen2017] | I / II | N=138 | 81kb | 38,589x | 6,182x | 59-71% |
| Breast, Lung, Prostate [Razavi2017b] | metastatic | N=124 | 2100kb | 60,000x | 3,000-4,000x | 89% |
| Non-ctDNA-based tests |  |  |  |  |  |  |
| Cancer Type | Stage | TP+FN / Total |  | Modality | FN Set | Clin. Sens. |
| Ovarian [Menon2009] | 48% I/II | 42+5 / 50,078 |  | CA-125 + ultrasound | 1yr followup | 89.4% |
| Colorectal [Imperiale2014] | 93% I/II/III | 60+5 / 9,989 |  | FIT + DNA | Colonoscopy | 92.3% |

Analysis of late-stage cancer patients, in whom ctDNA burden is higher [Bettegowda2014], offers a means to probe the limits of mutation detection. Razavi and colleagues presented data from 124 metastatic breast, lung, and prostate cancer patients using a sequencing panel covering 2.1 Mb at an average effective depth of 3,000-4,000x, achieving a clinical sensitivity of 89% [Razavi2017b]. While the 89% sensitivity achieved in metastatic patients is much higher than that achieved by similar sequencing depth (but smaller ROI) in early stage patients, it compares poorly to the performance of previously described early detection assays. Menon and colleagues described the prospective clinical validation of multimodal screening for ovarian cancer, combining measurement of serum CA-125 with ultrasound. The trial evaluated 50,078 individuals, of whom 47 had cancer within one year of trial completion (48% of cancers in stage I/II) [Menon2009]. The screening approach achieved 89.4% clinical sensitivity (42 TP detected with 5 FN discovered in 1-year followup). Imperiale and colleagues evaluated the performance of a fecal test for colorectal cancer, combining detection of occult blood with measurement of DNA markers in 9,989 individuals, achieving a clinical sensitivity of 92.3%, benchmarked relative to colonoscopy (which remains the gold standard for screening) [Imperiale2014]. Thus even given an unrealistic advantage (assuming ctDNA burdens equivalent to those present in metastatic cancer patients), the ctDNA-based mutation detection approach is unable to reach the sensitivity of existing assays for early cancer detection, making the exorbitant cost of sequencing difficult to reconcile.

## Conclusions

The advent of high-throughput sequencing, coupled with the demonstration that ctDNA can be detected non-invasively in plasma at various stages of malignancy, has led to significant investment in mutation-based ctDNA assays for early detection. However, there has yet to be a systematic exploration of the statistical and biological limits of such an approach. Here we have demonstrated that intrinsic biological characteristics likely set a performance bound for mutation detection that is insufficient to achieve high sensitivity and specificity in early-stage cancer detection using ctDNA detection alone.

The fundamental limitations described by the binomial model of tumor fraction pertain to *quantity* (because tumor-derived alleles are present at very low concentrations in bulk plasma, large volumes of plasma and huge sequencing bandwidth are required to detect even *one* tumor-derived molecule) and the specificity challenges posed by somatic heterogeneity arise from the requirement to interpreting a sample from a single patient at a single time, with no other context. Relaxation of either constraint may admit viable strategies for early detection.

**Box 1** summarizes a number of blood-based analytes with potential applicability for early cancer screening. While diverse, these analytes all have in common that they occupy more accessible points in the concentration-quantity space than ctDNA: options exist to find concentrated compartments of tumor-derived material, high-copy-number analytes (e.g., protein and RNA) that are present in measurable quantity even at low concentration. Particularly appealing opportunities arise in the search for signals that are not tumor-derived: if the tumor comprises 0.01% or less of circulating material, perhaps there is relevant signal in the other 99.99%, such as measurement of the body’s own immunosurveillance response to the presence of ‘foreign’ tumor cells. Integration of germline risk information may also boost the predictive power of a phenotype classifier [Shieh2016]. Computational integration of these multi-analyte signals may further provide improved power for phenotype classification [Argelaguet2017].

### Box 1

Biological components other than ctDNA with potential for cancer screening.

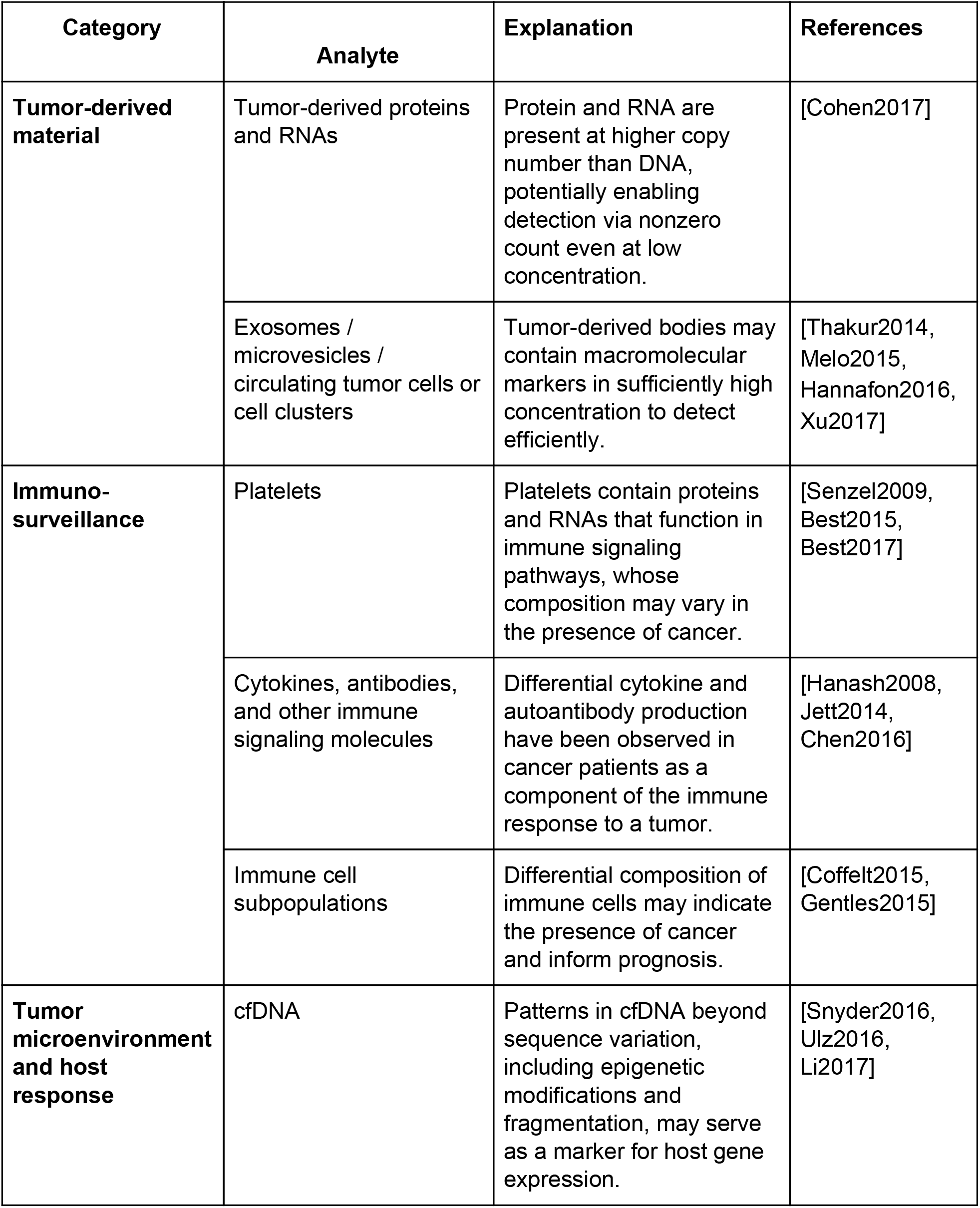

Furthermore, the screening context offers a unique opportunity to use longitudinal data on single individuals to improve accuracy. As cancer screening tests are typically repeated on a 1-3 year cycle, it may become possible to use not only an individual’s most recent test result but also his or her previous test results to precisely estimate changes in health status over time. Such a longitudinal testing approach may be able to use individualized data to reduce error by calibrating a predictor to individual baselines or known individual patterns of somatic variation and could also incorporate phenomena in the larger medical record. While such systems are current open areas of research, recent results on modeling of electronic medical record data suggest that the integration of structured data (such as diagnostic results) with temporal variables can improve accuracy in practice [Fiterau2017].

It has long been appreciated that early detection and intervention are perhaps the most effective means for reducing cancer related mortality. This significant unmet need has long motivated the search for accurate detectors of early-stage cancer, including potential signals in tumor-derived cell-free DNA. While statistical analysis suggests that sequencing of this ctDNA alone may be insufficient to achieve clinical sufficiency in this task, the presence of additional biological signals in the blood combined with multi-analyte or longitudinal analysis methods motivates further research towards the development of clinically usable early detection protocols.

## Acknowledgments

We thank Christina Curtis, Adam Drake, Kate Niehaus, Gabriel Otte, Girish Putcha, and David Weinberg for their suggestions and feedback on the manuscript.

## Supplemental Information

Attached is `make_figures.py,` a Python 2.7 script. It contains an implementation of the analytical binomial model as well as the reanalysis performed on the data from [Phallen2017], and is a self-contained package to fully regenerate figures 1 and 2.

